# First Metagenome-Assembled Genomes from the Historic Morrow Plots Reveal Management-Associated Dominance of Archaeal Ammonia Oxidizers

**DOI:** 10.64898/2026.03.24.714042

**Authors:** Viet D. Nguyen, Chushu Gao, Cory Gardner, Zhine Wang, Andrew J Margenot, Laibin Huang, Tae-Hyuk Ahn

**Author notes:** To whom correspondence should be addressed Laibin Huang, Tae-Hyuk Ahn.

## Abstract

Soil microbial communities underpin both soil health and agricultural productivity, yet genome-resolved resources from long-term field experiments remain limited. Here, we present a genome-resolved metagenomic dataset from the historic Morrow Plots long-term experiment in central USA, comprising 33 shotgun metagenomes collected across diverse crop rotation and fertilization treatments in year 149 of the experiment. Using a co-assembly, multi-binner workflow, we recovered 230 metagenome-assembled genomes (MAGs), including 44 archaeal and 186 bacterial genomes spanning multiple soil-associated phyla. Among these, 59 MAGs were linked to nitrogen-cycling functions, including ammonia- and nitrite-oxidizing lineages. The dataset also includes genome quality metrics, taxonomic classification, and treatment-resolved abundance patterns across different management regimes. Importantly, these nitrogen guild MAGs enable comparative analyses of nitrifier ecology, genome diversity, and functional variation linked to management in agricultural soils. Together, these resources establish a unique benchmark for studying how agricultural practices shape soil microbial communities at genome level, with associated long-term crop yield and soil fertility changes since the experiment’s inception in 1876.

## Background & Summary

Nitrogen cycling is one of the major determinants of agricultural productivity and sustainability because the interplay of mineralization, nitrification, assimilation, and loss pathways (e.g., denitrification) regulates both bioavailable nitrogen and environmentally relevant losses such as nitrate leaching and gaseous emissions^1-3^. Within these microbially-mediated transformation pathways, ammonia oxidation, the conversion of ammonia to nitrite, represents a key regulatory step in nitrification^4^. This process is primarily driven by ammonia-oxidizing archaea (AOA) and bacteria (AOB), as well as complete ammonia oxidizers (Comammox)^5^, with AOAs frequently dominating ammonia-oxidizer communities in soils^5-7^. Although marker-gene surveys and process-based studies have provided valuable insights into nitrifier ecology in soils, the genome-resolved ecological context that links long-term management histories to specific lineages and their metabolic potential remain limited^6,8,9^.

Genome-resolved metagenomics enables the reconstruction of metagenome-assembled genomes (MAGs), allowing taxonomic composition and functional potential to be examined together^6,10,11^ to help explain agroecosystem nitrogen processes. In soil systems, however, genome recovery remains challenging because of high community diversity, strain heterogeneity, and uneven abundance distributions^6,8^. Recent advances in co-assembly strategies, multi-binner approaches, consensus refinement, and standardized quality control have improved the recovery of soil MAGs^11-15^. Yet, relatively few genome-resolved datasets are available, in particular from long-term agricultural experiments where management practices have been applied consistently for more than a century^8,9^ and thus may better reflect at-equilibrium conditions of soil processes in agroecosystems.

The long-term Morrow Plots experiment at the University of Illinois Urbana–Champaign provides a valuable long-term experimental system for examining the effects of sustained crop rotation and fertilization regimes on soil microbial communities^16-18^. Established in 1876, the experiment contrasts monocropping system, like continuous corn (CCC), crop rotations including corn– soybean (CS) and corn–oat–alfalfa (COA), with each being managed without fertilization or fertilization with synthetic fertilizer or manure^16^. Additionally, we sampled a non-crop system, such as sod–grass (SG) for reference. The long duration and well-characterized management history make the Morrow Plots highly suited for generating reference datasets that offer a benchmark for comparative analysis of agricultural soil microbiomes, in particular in the US Midwest region of maize (*Zea mays* L.)-based agroecosystems responsible for a third of global maize and soybean (*Glycine max* L.) production.

Here, we present a genome-resolved metagenomic dataset generated from 33 soil metagenomes spanning 12 management combinations in the Morrow Plots. Figure 1 summarizes the experimental framework, sequencing workflow, and genome reconstruction strategy used to generate this resource, and Table 1 summarizes read filtering, co-assembly, and MAG recovery statistics across treatment groups. Through a standardized co-assembly and multi-step binning pipeline^9^, we reconstructed a catalog of 230 MAGs, including 44 archaeal and 186 bacterial genomes, together with associated genome quality statistics, taxonomic assignments, functional annotations, and treatment (crop rotation × fertility management)-level abundance information.

**Table 1.**
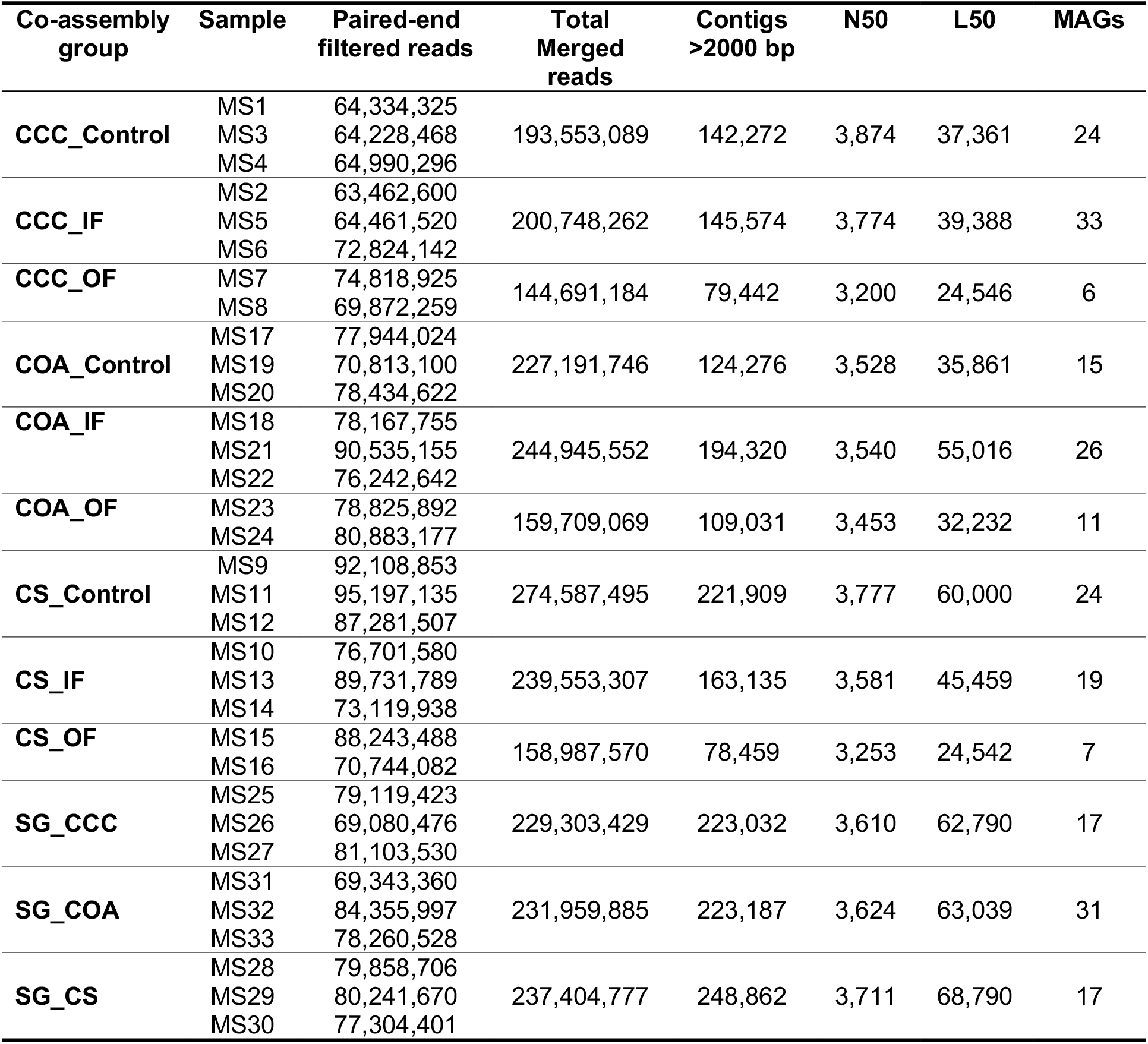
Summary of read filtering, co-assembly, and MAG recovery statistics for the Morrow Plots metagenomic dataset.

**Figure 1.**
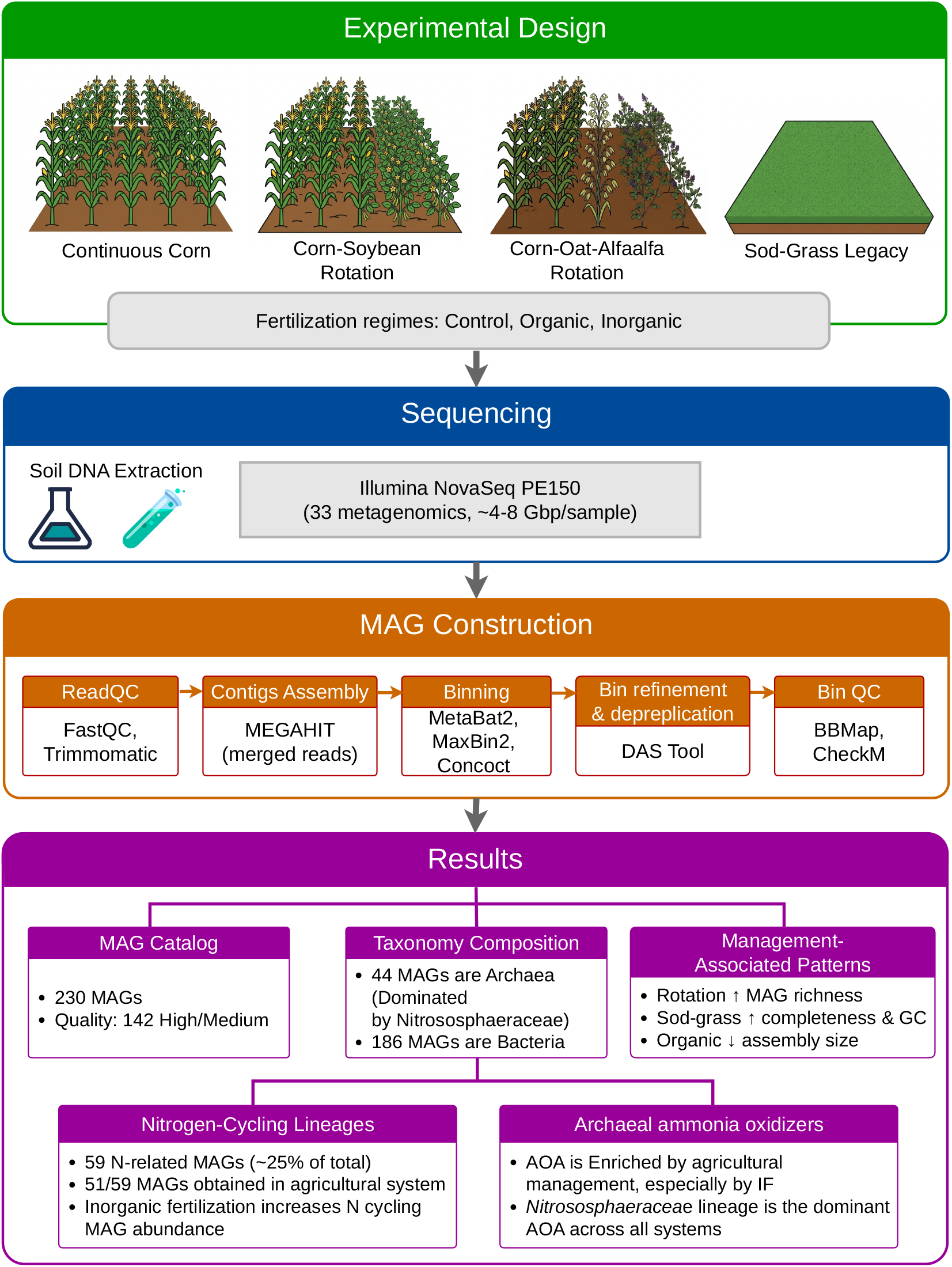
Experimental design, sequencing workflow, and overview of the genome-resolved metagenomic dataset generated for soils sampled in year 149 of the long-term Morrow Plots experiment in Illinois, USA.

Across the 33 soil metagenomes, the genome reconstruction workflow yielded a catalog that includes 1 high-quality MAG, 141 medium-quality MAGs, and 88 low-quality MAGs according to modified MIMAG criteria^11^. Figure 2 provides a descriptive overview of the reconstructed MAG collection, including phylogenetic placement, taxonomic composition, and selected genome characteristics across management groups. The catalog demonstrates a genome-resolved foundation for linking long-term management to microbial lineage composition and functional potential. Specifically, the interactive effects of fertilization and crop rotations on MAG richness were observed, with increased MAG richness under inorganic fertilization condition, particularly in continuous corn compared to corn rotated with other crops. Additionally, the sod–grass legacy condition exhibited distinct genome properties, including higher completeness and GC content. Organic fertilization was associated with smaller assembly size in workflow summaries, suggesting that management may influence both ecological composition and genome recoverability^6,8^.

**Figure 2.**
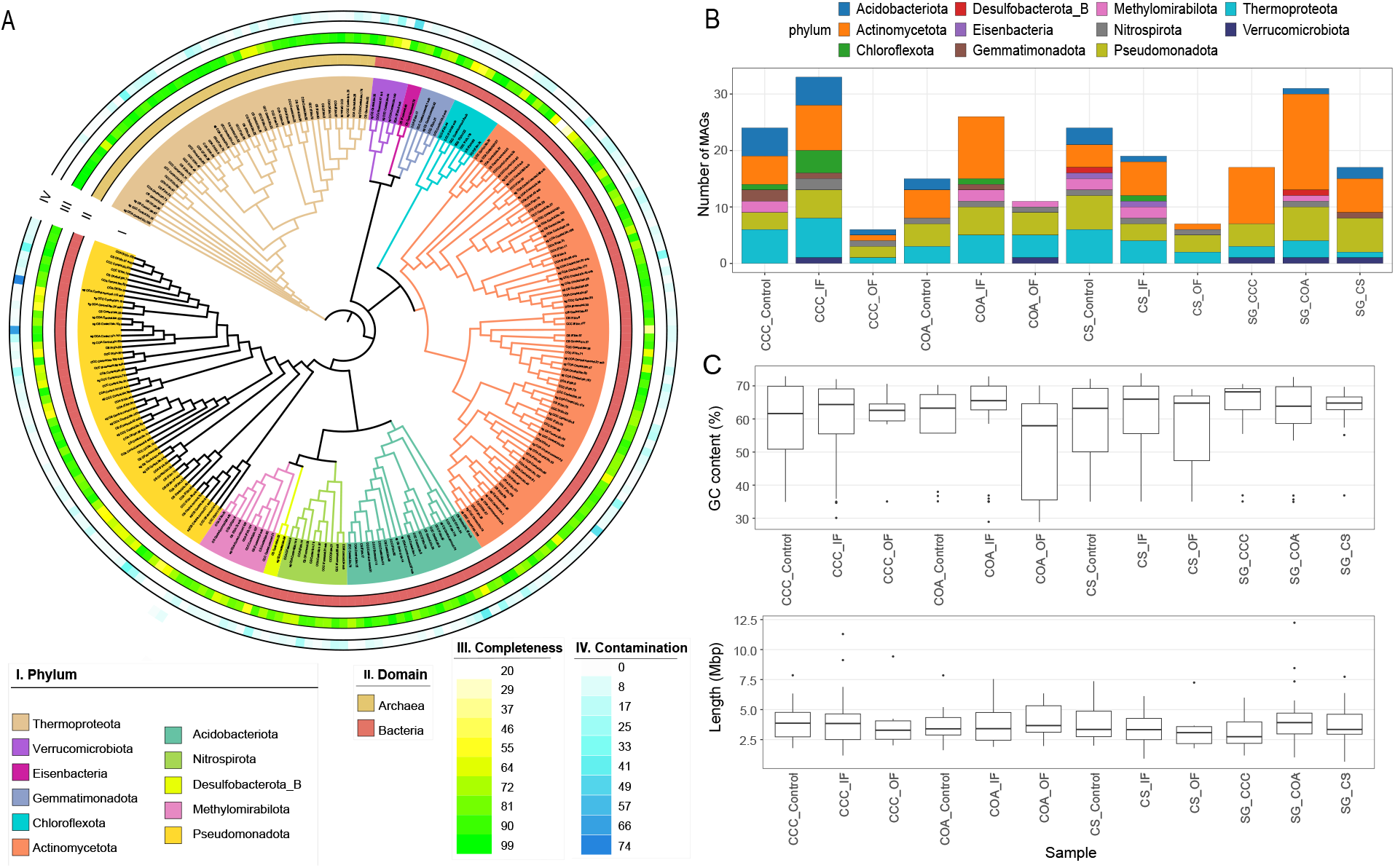
Phylogenetic placement, taxonomic composition, and genome characteristics of reconstructed MAGs. **(A)** Circular phylogenetic tree of 230 MAGs recovered from Morrow Plot topsoil, colored by phylum. Outer rings indicate domain (Archaea/Bacteria), genome completeness, and contamination. **(B)** Stacked bar plots showing the distribution of MAGs across phyla within each cropping system and fertilization regime. **(C)** Boxplots summarizing GC content and genome length (Mbp) across management groups, illustrating variation in genome properties among treatments.

To place the genome-resolved catalog in the context of broader community composition, we also compared taxonomic profiles derived from read-based classification with Kraken2^19^ to those inferred from reconstructed MAGs (Supplementary Figure S1). Read-based profiles showed high relative abundances of Actinomycetota, Pseudomonadota, Acidobacteriota, and Chloroflexota across treatments, consistent with typical agricultural soil communities, whereas MAG-derived profiles recovered many of the same dominant phyla though with notable differences in proportional representation. For instance, archaeal lineages affiliated with Thermoproteota (Nitrososphaeraceae) were proportionally more prominent in the MAG dataset relative to their representation in total reads, reflecting the successful targeted recovery of ammonia-oxidizing archaea (AOA) genomes (Figure 2A). Conversely, certain bacterial phyla abundant in the read-based dataset (e.g., Actinomycetota) were underrepresented among MAGs, likely due to the challenges of assembling genomes from highly diverse and strain-heterogeneous taxa (Supplementary Figure S1). Despite these biases, the MAG-based approach recovered several low-abundance phyla, including Nitrospirota, which provides a new avenue for studying complete nitrifier (Comammox) in agricultural soils (Supplementary Figure S2). Together, these differences highlighted expected assembly and binning biases in complex soil metagenomes while also demonstrating how genome-resolved analysis both complements and extends read-based community profiles.

Among the reconstructed genomes, 59 MAGs known to regulate nitrogen cycling in the soil represented approximately 25% of the total catalog. The majority of these nitrogen-cycling MAGs were recovered from cropping systems (51 of 59), providing a valuable resource for examining the effects of crop production on nitrogen-cycling populations across long-term management conditions. Within this subset, archaeal ammonia oxidizers affiliated with the family Nitrososphaeraceae were well represented with multiple genus-level clades detected across all treatments, including JAFAQB01, JARBAU01, TA-21, DASQQB01, TH1177, Nitrosocosmicus, and additional unclassified lineages (Figure 3). Their broad presence across management combinations, coupled with lineage-level phylogenetic resolution, supports the ecological prominence of AOA in these soils and extends prior marker-gene observations to a genome-resolved context^5-7,20^. In addition, Nitrospira-related genomes (Supplementary Figure S2), including comammox-associated lineages, were detected within the catalog, highlighting the representation of complete nitrification potential within long-term managed soils^21,22^ and further extending the functional scope of the dataset.

**Figure 3.**
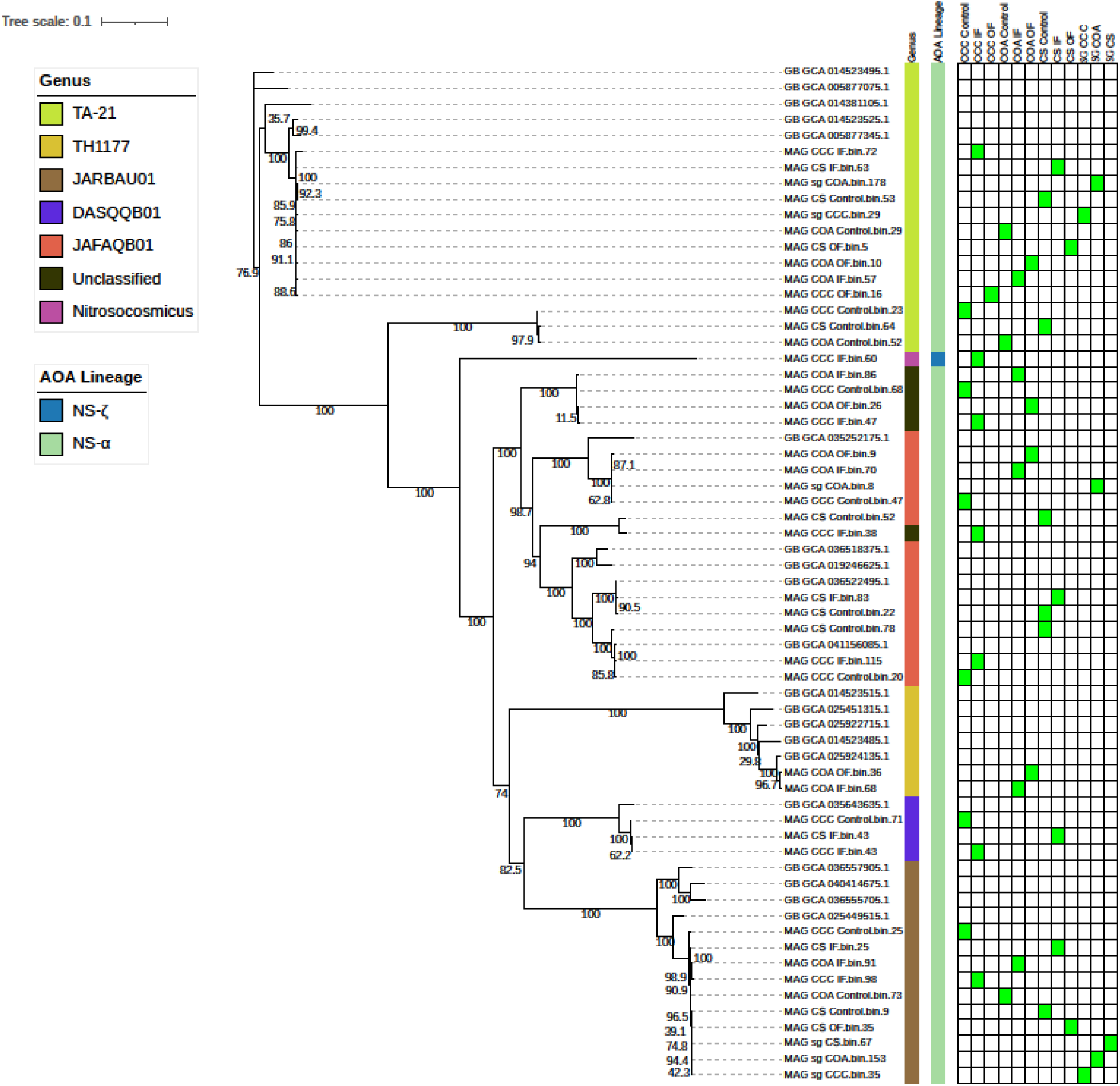
Phylogenetic placement and distribution of ammonia-oxidizing archaeal MAGs. Maximum-likelihood phylogenetic tree of archaeal MAGs affiliated with Nitrososphaeraceae recovered from Morrow Plot soils together with GTDB reference genomes from NCBI GenBank assembly accessions. Colored strips indicate genus-level GTDB classifications and broader AOA lineage assignments. The presence–absence heatmap shows detection of each MAG across cropping systems and fertilization treatments, with green cells indicating presence within a treatment group.

Collectively, this study provides a curated genome-resolved resource from one of the world’s longest-running agricultural experiments^16,17^. The dataset captures taxonomic diversity, genome quality metrics, and nitrogen-cycling potential across 149 years of sustained management, offering a foundation for comparative analyses of soil microbial evolution, nutrient cycling capacity, and agroecosystem sustainability.

## Methods

### Soil sampling and experimental design

Soil samples were collected from the historic Morrow Plots (est. 1876), the oldest continuous agricultural field in the United States, which are maintained by the University of Illinois Urbana-Champaign, IL^16,17^. The experiment site is situated on a fine, smectitic, mesic Aquic Argiudoll (Flanagan series), developed on loess parent material. The plots represent long-term cropping systems maintained for nearly 150 years, including continuous maize (CCC), a maize–soybean rotation (CS), a three-year maize–oat (*Avena sativa*)–alfalfa (*Medicago sativa*) rotation (COA) ^16^. In addition, sod-grass (SG) legacy plots were sampled as non-farming controls. Within each cropping system, soils were managed under unfertilized control (Control), inorganic fertilization (IF), or organic fertilization (OF) regimes (Figure 1), established in 1904. The SG forms the perimeter surrounding the replicated crop rotation × fertilization treatments. Established in 1904, the SG does not receive any nutrient inputs.

### DNA extraction and sequencing

Soil DNA was extracted from ~0.30 g of wet soil using the DNeasy Powersoil kits following the manufacturer’s instructions (Qiagen, CA, USA), and sequenced on an Illumina NovaSeq 6000 platform using 150 bp paired-end reads (PE150)^23^ at the Roy Carver Biotechnology Center at the University of Illinois at Urbana-Champaign. A total of 33 soil metagenomes were generated across these 12 treatments: CCC_Control, CCC_IF, CCC_OF, CS_Control, CS_IF, CS_OF, COA_Control, COA_IF, COA_OF, SG_CCC, SG_CS, and SG_COA. The sequencing depth ranged from approximately 4–8 Gbp per sample with a range of 64 to 95 million reads. Raw reads have been deposited in the NCBI Sequence Read Archive (SRA) under the BioProject accession listed in the Data Availability section.

### Metagenomic read processing, co-assembly and genome binning

For genome reconstruction, samples were grouped by replicates and co-assembled within each treatment. This design enabled treatment-level recovery of genomes while preserving the long-term experimental structure of the field system. Metagenomic read processing and genome reconstruction followed a standardized multi-step workflow^9^. In brief, raw reads were evaluated using FastQC v0.11.5^24^ and trimmed using Trimmomatic v0.36^25^ to remove reads shorter than 70 bp and bases with Phred scores below 30. Trimmed reads were then evaluated again using FastQC^24^, and MultiQC^26^ was used to summarize results.

To reduce method-specific bias in genome recovery, we applied a co-assembly and multi-binner strategy. For each treatment group, quality-filtered reads were co-assembled using MEGAHIT v1.2.9^27^. Only contigs generated by MEGAHIT with at least 2000 bp in length were retained for downstream binning. Assembly quality was assessed with QUAST^28^, and the filtered contigs were then binned using three complementary algorithms. MetaBAT2 v2.12.1^12^ was run with a minimum contig length of 2000 bp. MaxBin2 v2.2.7^13^ was executed with a probability threshold of 0.8 and minimum contig length of 2000 bp. CONCOCT v1.1.0^29^ was applied after cutting assemblies into 10 kbp fragments, and coverage profiles were generated by mapping reads to assemblies using BBMap v39.33^30^. Initial bin sets were refined using DAS Tool v1.1.7^31^ with the blastp search engine and score threshold of 0.6. This consensus refinement step generated a non-redundant set of metagenome-assembled genomes (MAGs) for quality assessment. MAG completeness and contamination were evaluated using CheckM v1.2.4^14^. Ribosomal RNA genes were identified using Barrnap v0.9^32^, and 5S, 16S, and 23S gene counts were parsed from GFF outputs. Transfer RNA genes were predicted using tRNAscan-SE v2.0.12^33^ with default search parameters, following the standards proposed by Bowers et al.^11^.

### Taxonomic annotation and genome quality characterization

Recovered MAGs were taxonomically classified using GTDB-Tk v2.5.2^15^ against database Release226. Taxonomic assignments were reported from domain to the lowest confidently assigned rank. Genome statistics including completeness, contamination, genome size, GC content, and marker gene content were compiled for all recovered MAGs (Table S1). These metrics were used to summarize the overall quality and diversity of the MAG catalog and were included in the deposited metadata tables. Archaeal MAGs were further evaluated in the context of ammonia-oxidizing lineages, particularly within Nitrososphaeraceae and Nitrospira, using phylogenetic placement relative to GTDB reference genomes (Figure 3). MAGs belonging to these groups were classified as nitrogen-associated genomes and retained for downstream analyses.

### Read mapping and treatment-level abundance profiling

To characterize taxonomic composition directly from metagenomic reads, paired-end sequencing reads were grouped by treatment and classified using Kraken2^19^ against the k2_standard_20230605_db reference database. For each treatment group, reads from all associated samples were pooled and analyzed together to generate treatment-level taxonomic profiles. Kraken2 reports were summarized at the phylum level and used to compare read-based community composition with the taxonomic distribution inferred from reconstructed MAGs.

### Phylogenetic analyses, visualization, and statistical summaries

Phylogenetic trees were generated separately for the full MAG catalog and the ammonia-oxidizing MAG subset. For the full catalog, representative marker-based phylogenetic placement was obtained using GTDB-Tk outputs and visualized as a circular tree with metadata rings indicating domain assignment, completeness, and contamination. For the archaeal subset, Nitrososphaeraceae-affiliated MAGs were analyzed together with GTDB reference genomes to generate a maximum-likelihood phylogenetic tree. Presence–absence information across treatment groups was incorporated as an accompanying heatmap.

Phylogenetic relationships among archaeal ammonia-oxidizing MAGs were inferred using conserved marker genes identified during GTDB-Tk^15^ classification. Marker gene alignments were used to construct a maximum-likelihood phylogenetic tree using IQ-TREE v3.0.1^34^, and branch support values were calculated using bootstrap analysis. The resulting tree was visualized and annotated using the Interactive Tree of Life (iTOL v6) ^35^. Lineage annotations were incorporated as color strips to distinguish archaeal ammonia-oxidizing clades (e.g., JAFAQB01, JARBAU01, TA-21, DASQQB01, TH1177, and Nitrosocosmicus-related lineages) and broader AOA lineage groupings. Presence–absence patterns across management treatments were visualized as heatmaps aligned to the phylogeny.

All statistical analyses and graphical visualizations were conducted in R 4.5.2 using custom scripts. Boxplots and genome feature comparisons were generated using ggplot2^36^, and stacked bar plots summarizing taxonomic distributions were created using ggplot2-based workflows^37^. Package versions and software details are provided in Supplementary Table S2.

## Data Records

Raw shotgun metagenomic reads from 33 soil samples have been deposited in the NCBI Sequence Read Archive under BioProject PRJNA1435883. The 230 reconstructed metagenome-assembled genomes (MAGs) are associated with NCBI BioSample accession numbers SAMN56535406–SAMN56535635. Processed datasets and supporting files are publicly available in Zenodo (https://doi.org/10.5281/zenodo.19119299), including compressed MAG files and tables containing sample metadata, MAG quality metrics, taxonomic assignments, nitrogen related annotations, treatment level abundance summaries, and co assembly statistics. Supplementary Table S1 provides detailed information for all reconstructed MAGs. Analysis scripts used for genome reconstruction, annotation, abundance estimation, and figure generation are available via GitHub (https://github.com/BioHPC/MorrowPlotsMAGs).

## Technical Validation

MAG quality was evaluated using CheckM v1.2.4^14^ with the lineage-specific workflow to estimate completeness and contamination based on single-copy marker genes. Genome statistics, including genome size, GC content, and contiguity metrics, were examined to assess structural integrity. Ribosomal RNA genes were identified using Barrnap v0.9^32^, and transfer RNA genes were predicted using tRNAscan-SE^33^ v2.0.12 to evaluate gene content completeness in accordance with MIMAG^11^ guidelines.

Following MIMAG^11^ standards, high-quality MAGs were defined as ≥90% completeness and ≤5% contamination with the presence of 5S, 16S, and 23S rRNA genes and ≥18 tRNAs. Medium-quality MAGs were defined using a modified threshold of >70% completeness and <10% contamination to ensure reliable genome representation while accounting for the inherent fragmentation typical of soil assemblies. 142 MAGs were classified as medium or high quality, supporting the reliability of the genome-resolved dataset for downstream ecological analyses.

## Data Availability

Raw sequencing data are available in the NCBI Sequence Read Archive under BioProject PRJNA1435883. The 230 reconstructed metagenome assembled genomes (MAGs) are associated with NCBI BioSample accession numbers SAMN56535406–SAMN56535635. Processed datasets and supporting files are available in Zenodo (https://doi.org/10.5281/zenodo.19119299), including compressed MAG files and tabular datasets containing sample metadata and result summaries.

## Code Availability

All scripts used in this study are available at https://github.com/BioHPC/MorrowPlotsMAGs, with an archived release deposited in Zenodo (https://doi.org/10.5281/zenodo.19119299).

## Author Contributions

T.-H.A. and L.H. conceived and supervised the study. L.H. contributed to experimental design, soil collection, funds for sequencing, biological interpretation, and manuscript revision. A.M. contributed to the experimental design, field management, soil collection, and manuscript revision. V.D.N. performed data processing, metagenome assembly, genome reconstruction, annotation, figure preparation, and drafted the manuscript. C.S.G. and Z.W. collected the soil sample and did the DNA extraction and sent the DNA for sequencing. T.-H.A. and C.G. generated the GitHub and Zenodo, and did manuscript revision. All authors reviewed and approved the final manuscript.

## Acknowledgements

We thank Ebun Omole-Ohonsi, Birch Fabregas, Sabita Ghimire for assistance with sample handling, sequencing coordination, data management, or helpful discussion. We also acknowledge the Morrow Plots program and the University of Illinois Urbana–Champaign for maintaining the long-term experimental infrastructure that made this dataset possible.

## References

1 Huang, L. Molecular and dual-isotopic profiling of the microbial controls on nitrogen leaching in agricultural soils under managed aquifer recharge. Environ. Sci. Technol. 57 (2023). 10.1021/acs.est.3c01356

2 Levintal, E. Nitrogen fate during agricultural managed aquifer recharge: Linking plant response, hydrologic, and geochemical processes. Sci. Total Environ. 864 (2023). 10.1016/j.scitotenv.2022.161206

3 Zhang, H., Xu, Y. & Kanyerere, T. A review of the managed aquifer recharge: Historical development, current situation and perspectives. Phys. Chem. Earth 118 (2020). 10.1016/j.pce.2020.102887

4 Kuypers, M. M. M., Marchant, H. K. & Kartal, B. The microbial nitrogen-cycling network. Nat Rev Microbiol 16, 263–276 (2018). 10.1038/nrmicro.2018.9

5 Huang, L. et al. Ammonia-oxidizing archaea are integral to nitrogen cycling in a highly fertile agricultural soil. ISME Commun 1, 19 (2021). 10.1038/s43705-021-00020-4

6 Lemos, L. N., Mendes, L. W., Baldrian, P. & Pylro, V. S. Genome-resolved metagenomics is essential for unlocking the microbial black box of the soil. Trends Microbiol. 29 (2021). 10.1016/j.tim.2021.01.013

7 Leininger, S. et al. Archaea predominate among ammonia-oxidizing prokaryotes in soils. Nature 442, 806–809 (2006). 10.1038/nature04983

8 Anthony, W. E. From soil to sequence: filling the critical gap in genome-resolved metagenomics is essential to the future of soil microbial ecology. Environ. Microbiome 19 (2024). 10.1186/s40793-024-00599-w

9 Brandão Gontijo, J. et al. Depth-dependent Metagenome-Assembled Genomes of Agricultural Soils under Managed Aquifer Recharge. Scientific Data 12 (2025). 10.1038/s41597-025-05218-y

10 Parks, D. H. Recovery of nearly 8,000 metagenome-assembled genomes substantially expands the tree of life. Nat. Microbiol. 2 (2017). 10.1038/s41564-017-0012-7

11 Bowers, R. M. Minimum information about a single amplified genome (MISAG) and a metagenome-assembled genome (MIMAG) of bacteria and archaea. Nat. Biotechnol. 35 (2017). 10.1038/nbt.3893

12 Kang, D. D. MetaBAT2: An adaptive binning algorithm for robust and efficient genome reconstruction from metagenome assemblies. PeerJ 2019 (2019). 10.7717/peerj.7359

13 Wu, Y. W., Simmons, B. A. & Singer, S. W. MaxBin 2.0: An automated binning algorithm to recover genomes from multiple metagenomic datasets. Bioinformatics 32 (2016). 10.1093/bioinformatics/btv638

14 Parks, D. H. CheckM: Assessing the quality of microbial genomes recovered from isolates, single cells, and metagenomes. Genome Res. 25 (2015). 10.1101/gr.186072.114

15 Chaumeil, P. A., Mussig, A. J., Hugenholtz, P. & Parks, D. H. GTDB-Tk: A toolkit to classify genomes with the genome taxonomy database. Bioinformatics 36 (2020). 10.1093/bioinformatics/btz848

16 Raglin, S. S., Soman, C., Ma, Y. & Kent, A. D. Long Term Influence of Fertility and Rotation on Soil Nitrification Potential and Nitrifier Communities. Frontiers in Soil Science Volume 2 - 2022 (2022). 10.3389/fsoil.2022.838497

17 Caldrone, S. L., Margenot, A. J. & Morrow Plots Data Curation Working, G. From complex histories to cohesive data, a long-term agricultural dataset from the Morrow Plots. Sci Data 11, 1145 (2024). 10.1038/s41597-024-03984-9

18 Soman, C., Li, D., Wander, M. M. & Kent, A. D. Long-term fertilizer and crop-rotation treatments differentially affect soil bacterial community structure. Plant and Soil 413, 145–159 (2017). 10.1007/s11104-016-3083-y

19 Wood, D. E., Lu, J. & Langmead, B. Improved metagenomic analysis with Kraken 2. Genome Biol 20, 257 (2019). 10.1186/s13059-019-1891-0

20 Prosser, J. I. & Nicol, G. W. Archaeal and bacterial ammonia-oxidisers in soil: the quest for niche specialisation and differentiation. Trends in Microbiology 20, 523–531 (2012). 10.1016/j.tim.2012.08.001

21 Daims, H. et al. Complete nitrification by Nitrospira bacteria. Nature 528, 504–509 (2015). 10.1038/nature16461

22 van Kessel, M. A. H. J. et al. Complete nitrification by a single microorganism. Nature 528, 555–559 (2015). 10.1038/nature16459

23 Modi, A., Vai, S., Caramelli, D. & Lari, M. In Bacterial Pangenomics: Methods and Protocols (eds Alessio Mengoni, Giovanni Bacci, & Marco Fondi) 15–42 (Springer US, 2021).

24 Andrews, S. FastQC: A quality control tool for high throughput sequence data. Available online at: http://www.bioinformatics.babraham.ac.uk/projects/fastqc/ (2010).

25 Bolger, A. M., Lohse, M. & Usadel, B. Trimmomatic: A flexible trimmer for Illumina sequence data. Bioinformatics 30 (2014). 10.1093/bioinformatics/btu170

26 Ewels, P., Magnusson, M., Lundin, S. & Kaller, M. MultiQC: summarize analysis results for multiple tools and samples in a single report. Bioinformatics 32, 3047–3048 (2016). 10.1093/bioinformatics/btw354

27 Li, D., Liu, C. M., Luo, R., Sadakane, K. & Lam, T. W. MEGAHIT: An ultra-fast singlenode solution for large and complex metagenomics assembly via succinct de Bruijn graph. Bioinformatics 31 (2015). 10.1093/bioinformatics/btv033

28 Gurevich, A., Saveliev, V., Vyahhi, N. & Tesler, G. QUAST: quality assessment tool for genome assemblies. Bioinformatics 29, 1072–1075 (2013). 10.1093/bioinformatics/btt086

29 Alneberg, J. Binning metagenomic contigs by coverage composition. Nat. Meth. 11 (2014). 10.1038/nmeth.3103

30 Bushnell, B. BBMap: A Fast, Accurate, Splice-Aware Aligner. (Ernest Orlando Lawrence Berkeley National Laboratory, Berkeley, CA (US), United States, 2014).

31 Sieber, C. M. K. Recovery of genomes from metagenomes via a dereplication, aggregation and scoring strategy. Nat. Microbiol. 3 (2018). 10.1038/s41564-018-0171-1

32 Seemann, T. barrnap 0.9: rapid ribosomal RNA prediction. Google Scholar 792 (2013).

33 Chan, P. P., Lin, B. Y., Mak, A. J. & Lowe, T. M. tRNAscan-SE 2.0: improved detection and functional classification of transfer RNA genes. Nucleic Acids Res 49, 9077–9096 (2021). 10.1093/nar/gkab688

34 Nguyen, L. T., Schmidt, H. A., von Haeseler, A. & Minh, B. Q. IQ-TREE: a fast and effective stochastic algorithm for estimating maximum-likelihood phylogenies. Mol Biol Evol 32, 268–274 (2015). 10.1093/molbev/msu300

35 Letunic, I. & Bork, P. Interactive Tree of Life (iTOL) v6: recent updates to the phylogenetic tree display and annotation tool. Nucleic Acids Res 52, W78–W82 (2024). 10.1093/nar/gkae268

36 Wickham, H. ggplot2: Elegant Graphics for Data Analysis. (Springer Publishing Company, Incorporated, 2016).

37 Kassambara, A. ggpubr: ggplot2-Based Publication Ready Plots. R package version 0.6.0 (2023).

