## Supplementary figures and images for "First Metagenome-Assembled Genomes from the Historic Morrow Plots Reveal Management-Associated Dominance of Archaeal Ammonia Oxidizers" (from another version of this preprint: version 1, 2026-03-25)

### Supplemental Figure 1

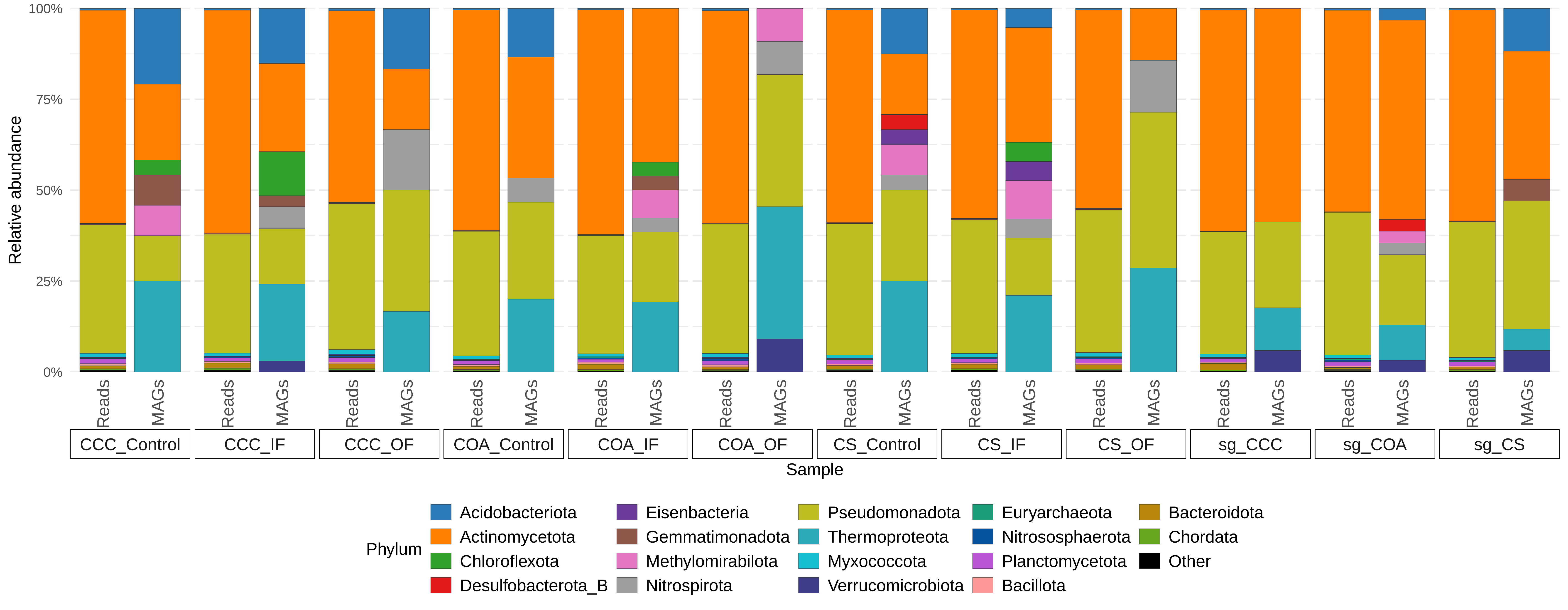

### Supplemental Figure 2

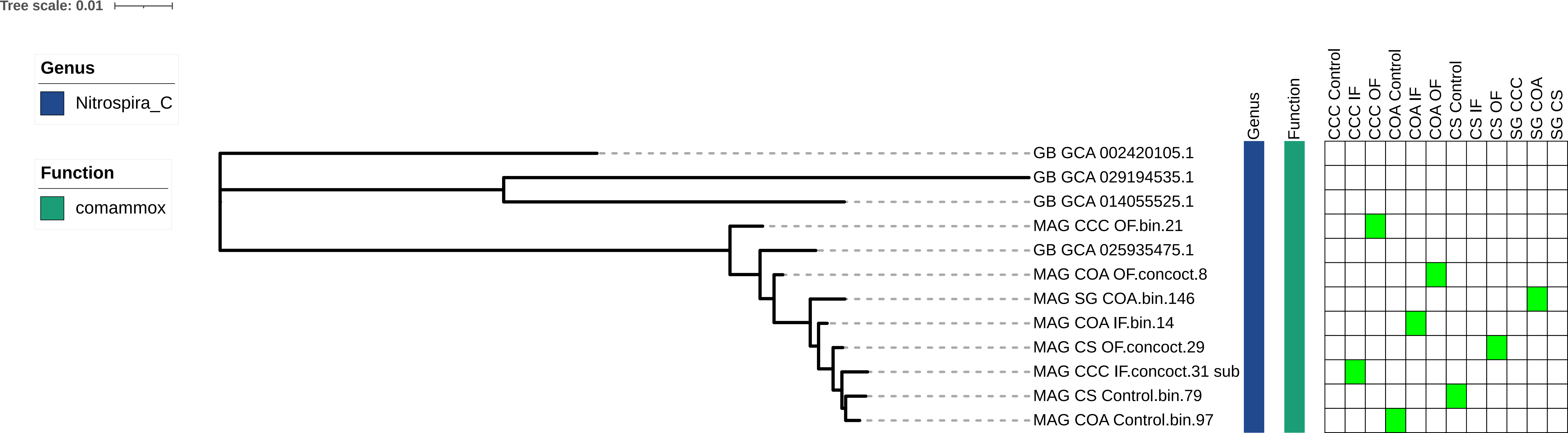
